# Depth-Sensitive Optical Property Characterization Using Multi-Frequency Laparoscopic SFDI

**DOI:** 10.64898/2026.02.04.703750

**Authors:** Elias Kluiszo, Luigi Belcastro, Rasel Ahmmed, Ulas Sunar

## Abstract

Accurate knowledge of tissue absorption (*μ*_*a*_) and reduced scattering 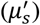 parameters are required to plan and monitor laparoscopic chemophototherapy (CPT) in ovarian cancer, including light dosimetry and quantitative fluorescence mapping of porphyrin-phospholipid (PoP) photobleaching and light-triggered doxorubicin (Dox) release. We implemented a depth-sensitive, multi-frequency laparoscopic spatial frequency domain imaging (SFDI) framework to improve optical-property estimation in layered tissue. A DMD-based laparoscope imaged two-layer phantoms with controlled optical contrasts and superficial thicknesses. Spatial-frequency subsets associated with different penetration depths were independently fit to recover *μ*_*a*_ and 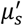, and compared with a two-layer diffusion model. Recovered 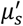 values remained bounded by the known layer references and shifted monotonically toward the superficial value as spatial frequency and top-layer thickness increased, approaching a single-layer response at high frequency/thick layers. Quantitative model comparison showed δ-P1 variants outperformed the standard diffusion approximation, reducing RMSPE between modeled and measured 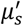 to 0.8-6.5% (silicone/silicone) and 1.6-8.3% (silicone/intralipid), whereas SDA errors reached ∼13.8% and 21.1%, respectively. This approach demonstrates multi-frequency laparoscopic SFDI as a practical initial step for depth-sensitive fluorescence correction for individualized CPT treatment planning and monitoring.

## 1. Introduction

Ovarian cancer is one of the most lethal gynecologic malignancies, in part because early-stage disease is often asymptomatic and current screening methods lack sufficient sensitivity and specificity for early detection [1,2]. Clinical assessment typically relies on transvaginal ultrasound (TVU), physical examination, and serum biomarkers such as cancer antigen 125 (CA125), but these approaches can yield false positives and have limited performance in detecting small-volume disease [2]. More invasive approaches, including endoscopic sampling and histopathology, can improve diagnostic accuracy but increase risk and patient discomfort. There is a strong need for minimally invasive, real-time intraoperative guidance that can improve tumor detection and imaging-guided drug delivery during ovarian cancer surgery.

Optical imaging techniques offer non-ionizing, high-sensitivity contrast and can be deployed intraoperatively. Spatial frequency domain imaging (SFDI) projects sinusoidal illumination patterns to quantify tissue absorption (*μ*_*a*_) and reduced scattering 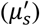 parameters that can change with pathological alterations in tissue structure and function [3–7]. SFDI has been applied in vivo and ex vivo across multiple disease sites, including breast, skin, and oral cancer [5,8,9]. Beyond marker-free contrast, SFDI can also improve quantitative fluorescence imaging by providing optical-property-based corrections that account for tissue-dependent attenuation of excitation and emission light, which is critical for applications such as photodynamic therapy (PDT) guidance [10–13].

Conventional SFDI is typically performed in open-field geometries and on relatively accessible tissue surfaces. Laparoscopy enables minimally invasive access to the abdominal and pelvic cavities through small incisions, providing real-time visualization during surgery. By coupling a modulated light source (e.g., a digital micromirror device, DMD) and a camera to an endoscope via fiber optics, SFDI can be translated to the laparoscopic setting for in vivo mapping of optical properties [14,15]. Laparoscopic SFDI is therefore promising for epithelial cancers in the peritoneal cavity, including ovarian cancer, where superficial and sub-surface lesions may be difficult to visualize with standard white-light imaging. In addition to wide-field coverage, SFDI offers tunable depth sensitivity through the spatial frequency (*f*_*x*_) of the projected patterns [16]. Higher spatial frequencies attenuate more rapidly in scattering media, reducing the effective sampling depth and increasing sensitivity to superficial layers, whereas lower spatial frequencies probe deeper tissue volumes. Belcastro et al. introduced a multi-frequency (multi-depth) processing framework that groups SFDI measurements into frequency subsets to estimate depth-dependent scattering contrast in layered tissue [17,18].

Wide-field layered-tissue analysis has also been explored for applications such as wound assessment and melanoma resection [19–21]. Extending depth-resolved SFDI to laparoscopy could enable assessment of heterogeneous tissue architectures, including discrimination between superficial epithelium (hundreds of micrometers thick) and underlying connective tissue surrounding tumors. A laparoscopic approach to depth-sensitive SFDI is particularly suited to resection surgeries and imaging-guided therapy within the abdominal cavity where peritoneal formations like omenta, mesenteries and ligaments are often the initial attachment site for disseminated tumors from ovarian and gastric cancers [22,23].

Depth-sensitive optical property estimation is also important for quantitative fluorescence imaging in addition to providing label-free contrasts. Endoscopic SFDI has been used to correct fluorescence signals for tissue absorption and scattering to quantify fluorophore concentrations in vivo [24,25]. Most fluorescence-correction models assume homogeneous optical properties and apply a single *μ*_*a*_ and 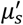 across the imaged volume. Depth sensitivity can enable more accurate fluorescence attenuation correction by weighting the optical-property contributions of superficial and deeper layers. This capability is particularly relevant for fluorescence-guided, light-activated drug delivery in chemophototherapy (CPT), where photodynamic therapy (PDT) is combined with chemotherapy (e.g., porphyrin-phospholipid, PoP, and liposomal doxorubicin, Dox) to trigger localized drug release and treatment response. Quantitative CPT monitoring requires the knowledge of *μ*_*a*_ and 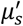 at both excitation and emission wavelengths to recover absolute drug fluorescence concentrations and to predict light penetration that governs drug activation.

Here, we apply multi-frequency processing to a DMD-based laparoscopic SFDI system previously developed for combined SFDI and quantitative fluorescence imaging [15]. Two-layer tissue-mimicking phantoms with controlled optical properties and variable superficial-layer thickness were imaged and processed using the multi-frequency approach described by Belcastro et al. [18]. The depth-dependent trends in recovered 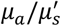 were compared against diffusion-based two-layer light-transport models, including cases with refractive-index mismatch, and additional measurements were performed on phantoms with scattering heterogeneity in depth to demonstrate depth sensitivity in a visually salient geometry. This study provides a step toward laparoscopy, enabling real-time, depth-resolved feedback for CPT and ensuring that light-triggered chemotherapy is delivered with unprecedented precision.

## 2. Materials and methods

### 2.1 Laparoscopic SFDI system

The laparoscopic SFDI system used in this study integrates the following components: a low noise research-grade EMCCD camera (Luca R, Andor, Belfast, Ireland), a modified Digital Micromirror Device (DLP4500, Texas Instruments, TX, USA) with its internal light source removed, three to four high power LEDs (High Power Collimated sources, Mightex, Ontario, Canada) with their respective LED controller (Universal 4-channel LED controller, Mightex, Ontario, Canada). The LEDs can be easily substituted, and their wavelengths are chosen according to the application. The LED output is combined using dichroic mirrors (Multi-Wavelength Beam Combiners, Mightex, Ontario, Canada) and is coupled to a liquid lightguide (5mm core, Mightex, Ontario, Canada) with a lightguide collimator (Mightex, Ontario, Canada). The lightguide transmits the LED output into the modified DMD, which is used to generate sinusoidal patterns and modulate the light intensity. Additionally, a manual filter holder (ThorLabs, NJ, USA) coupled with the LED combiner for precise fluorescence excitation.

The DMD output is then coupled with an imaging optical fiber, which is inserted in a laparoscope (8912.43, R. Wolf, Vernon Hills, IL, USA) and used to project the light patterns from its tip. A second imaging optical fiber in the laparoscope captures the reflected light and is coupled with the EMCCD camera for detection. Additional components in the detector includes a zoom coupler (Accu-Beam, TTI Medical, CA, USA), two 25 mm achromatic lenses, a filter wheel, and an aperture (ThorLabs, NJ, USA) before reaching the EMCCD camera. For the SFDI measurements, cross-polarizers were built into the front of the laparoscope’s distal end to reduce spectral reflection. A structural frame acted as a mechanical stop to ensure that the object surface was no closer than the minimal effective working distance of the scope [3,4,7,26]. Due to the divergence of light from the objective lenses, the frequency of the projected SFDI patterns will increase with distance, so the structural frame ensures that the target is always close to the optimal working distance. The camera was focused over the entire area of the projected SFDI pattern at a 4.0 × 4.0 cm field of view (FOV), which can be suitable in laparoscopic imaging during surgery and treatment planning.

### 2.2 Phantom models

To study the effectiveness of the laparoscopic SFDI system in detecting contrast in thin layers, tissue simulating phantoms with controlled optical properties have been manufactured. Two types of phantoms have been employed for this purpose: liquid phantoms made from Intralipid [27] and India ink [28], and silicone phantoms incorporating TiO2 particles and ink or graphite powder [29–31]. Intralipid phantoms have also been used for calibrating the instrument, as their absorption and scattering could be independently characterized by means of collimated transmission measurements using a wide-spectrum halogen light source and a spectrophotometer (Ocean Optics, Orlando, FL, USA).

Two multi-layer phantom models were implemented: one with both top and bottom layers made from silicone (silicone/silicone model) [32], and the other with a top layer of silicone and a bottom layer of intralipid (silicone/intralipid model). The silicone/silicone model is intended to validate the layer contrast of the endoscopic system at varying layer thicknesses. The silicone/intralipid model is intended to investigate changes in scattering behavior when there is a heterogenous index of refraction, a documented characteristic of solid tumors [33]. An increase in refractive index within cancerous regions is expected due to increased cell density and greater protein content within the cytoplasm, and differentiation between healthy and cancerous cells based upon refractive index analysis has been documented [34–36]. The index of refraction difference between silicone and intralipid (∼0.1) is of biological relevance, being similar to the difference between muscular and adipose tissue [37].

India ink (Speedball, Super Black, Statesville, North Carolina, USA) was selected as the absorbing agent due to its relatively uniform absorption spectrum over the entire measured wavelength range. Since the primary goal of this study is to model light scattering, it was important to minimize the spectral features introduced by absorbers, which could interfere with the scattering signal through spectral crosstalk during data analysis. The ink was first blended into a large volume of unpolymerized silicone at a concentration of 50 *μ*L/100 mL, and then distributed into separate batches for phantom fabrication. At this concentration, the absorption coefficient was expected to be approximately 0.015 mm^−1^ and was independently confirmed using optical measurements on a large phantom made from the solution to validate the phantom preparation method and to account for any unexpected deviations.

TiO2 of less than 25 nm particle size (Millipore Sigma, Burlington, MA, USA) was added to the uncured silicone mixture at a concentration of 2000 mg/100mL as a scattering agent and was dispersed via sonication to reduce particle aggregation that might affect particle distribution. The curing agent was then mixed with the unpolymerized silicone in a 1:10 ratio. To eliminate air bubbles formed during mixing, the blend was degassed in a vacuum chamber.

From each batch, five thin phantoms were fabricated for use as the top layers of the model, ranging in thickness from about 0.3 to 1.2 mm. The small quantities (0.5 to 3 ml) of the silicone mixture were dispensed using a syringe into 5 cm diameter petri dishes and allowed to spread evenly on a level surface. For the thinnest phantoms, a vortexer was used to vibrate the dish and encourage even distribution. Thickness measurements were taken using a micrometer at different locations (N = 10), with the mean and standard deviation values reported in Table 1. Remaining silicone from the batch was used to create a thicker, homogeneous phantom approximately 20-30 mm in depth for optical property quantification of the final batch (as mentioned prior).

**Table 1.**
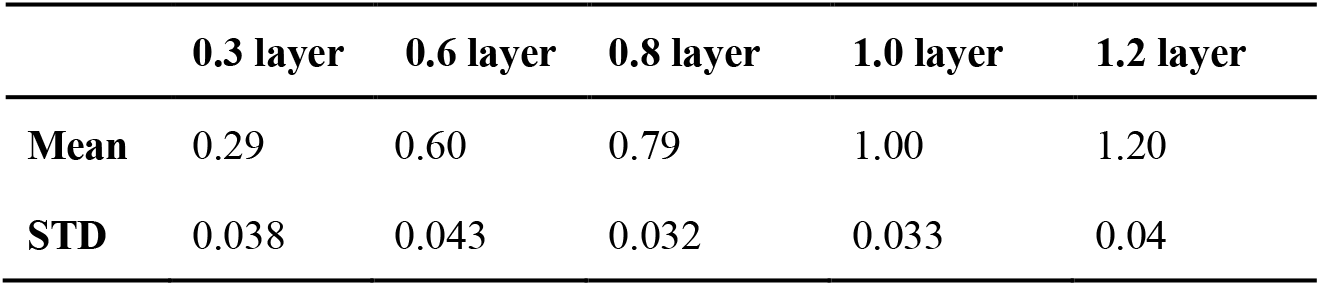
Measured thicknesses (mm) of superficial silicone layers used in two-layer phantoms (n = 10 each).

After generation of the thin, top phantom layers, a 35mm deep phantom base layer was created in a 3D printed mold to represent the semi-infinite bottom layer contribution within the two-layer model. TiO2 powder was added at a concentration of 500 mg/100mL and the ink at a concentration of 200 *μ*L/100 mL. Lastly, a mold of a cylindrical-conical shape was used to create a silicone phantom with a TiO2 concentration of 1000 mg/100mL and an India ink concentration of 50 *μ*L/100 mL. When submerged in an intralipid phantom, the conical shape of this phantom allows for differing surface areas of contributing 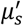 at different depths, allowing for a visual representation of how scattering contributions change at differing projected frequencies. In Fig. 9b, the cone was placed in a cup of the same height and filled to the brim with intralipid.

Liquid phantoms were created using 20% intralipid for scattering and the same India Ink for absorption. In the silicone/intralipid model, the same top layers were used and the intralipid bottom layer was made to the same 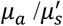 as the silicone bottom layer. A hollow, 35mm deep cylinder with a diameter slightly less than that of the top layer phantoms was 3D printed to contain the intralipid bottom layer while allowing the silicone top layer to be placed on top. Optical properties of all phantoms can be summarized in Table 2.

**Table 2:** Optical properties (*μ*_*a*_ and 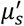 at 656 nm) of each phantom component used in layered models.

|  | Silicone/Silicone |  | Silicone/Intralipid |  | Conical |  |
| --- | --- | --- | --- | --- | --- | --- |
|  | Top | Bottom | Top | Bottom | Cone | Liquid |
| $\mu'_s$ (mm<sup>-1</sup>) | 3.11 | 0.83 | 3.11 | 0.81 | 1.43 | 2.06 |
| $\mu_a$ (mm<sup>-1</sup>) | 0.015 | 0.075 | 0.015 | 0.072 | 0.029 | 0.044 |

### 2.3 Image acquisition and processing

Following the characterization of all silicone and intralipid phantom batches, imaging data was collected on bi-layer phantoms using the previously described laparoscopic SFDI system. These two-layer configurations were made by placing the thin phantoms atop the semi-infinite homogeneous phantom, either silicone or intralipid, ensuring a contrast in scattering properties between layers. To reduce air gaps between both models, the top layer was carefully applied, and in the silicone/intralipid model, the intralipid base was overfilled slightly prior to top layer placement, and the silicone layer was placed on the convex meniscus to ensure full contact. For each acquisition, 21 spatial frequencies (*f*_*x*_) were captured, ranging from 0 mm^−1^ (uniform illumination) to 0.34 mm^−1^ in increments of 0.02 mm^−1^. These sinusoidal patterns were generated in MATLAB and projection equipment was adjusted to ensure the correct number of line pairs per mm. Using the traditional SFDI approach, with three phase projections for each spatial frequency, a total of 63 images were captured per phantom setup. All were made using a 656nm high power LED (High Power Collimated sources, Mightex, Ontario, Canada) sent through the DMD, with the EMCCD camera set to 250 ms exposure time at 4×4 camera binning. A dark image with no incoming signal was captured by the camera and subtracted from the acquired images to reduce the effect of noise on the acquired data.

The multi-frequency processing strategy implemented divides the full dataset into averaged subgroups of three spatial frequencies, each arranged in ascending order: Frequency averages did not repeat any frequency sections, so each average is derived from unique values only. Each frequency subset was independently analyzed to extract the absorption and reduced scattering coefficients, which are associated with distinct penetration depths as discussed in Section 1. The average spatial frequency of each subset is used in fluence modeling as described in the introduction. Data was analyzed under the assumption of a homogeneous medium, following the method described by Cuccia et al. [3], where the modulated signal amplitudes are first extracted and then calibrated using a homogeneous phantom with known optical parameters to derive diffuse reflectance. An iterative white Monte Carlo model is then used, adjusting input parameters 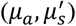 through optimization until the simulated diffuse reflectance aligns with the measured data, ultimately yielding the optical properties of the target sample. In future work, these depth-sensitive 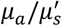 estimates will be used to implement depth-weighted fluorescence attenuation correction for drug fluorescence quantification during CPT.

### 2.4 Analytical Model

The data acquired from these models was plotted against the models of light fluence developed by Belcastro et al.[18]. Briefly, the contributions of each layer on the light fluence detected from the two-layer model can be ascribed to complementary weighting coefficients proportional to the actual layer thickness, which are then multiplied by each layer’s individual 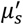, summing to the total 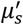 of the two-layer model.

The partial contributions of each layer along with their weighting coefficient on the 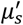 of the entire model is described in equation 1:

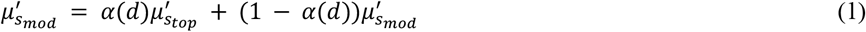

This weighting coefficient α as a function of layer depth can be acquired using the integral of the fluence from light passing through both layers and dividing the integral of the fluence from the top layer by the integral through the entire two-layer model.

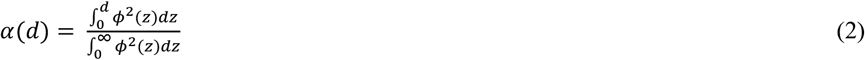

After acquiring the optical properties at each projected frequency, the fluence can be estimated using various analytical models, in this case we used the standard diffusion approximation (SDA) [38], the δ-P1 approximation [39,40], and a modified δ-P1 approximation developed by Belcastro et al.[18]. Applying these models to the acquired data serves as a method of validating the laparoscope’s sensitivity and as an investigation into potential solutions to the inverse problem, i.e. determining layers based upon derived optical properties or fluence alone.

To more accurately model the boundary conditions at the silicone/intralipid index of refraction mismatch, a weighted index of refraction term was used to derive the effective reflection coefficient (*R*,_*eff*_) in all three models:

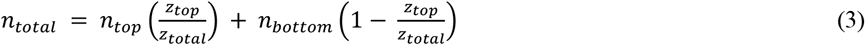

Where *z*_*top*_ represents the distance required to integrate *ϕ*^2^ up to *d*, and *z*_bottom_ represents the entire distance to integrate *ϕ*^2^. The weighted *n*_*total*_ value is used to calculate the effective reflection coefficient, which is then used to derive a constant determined by the choice of a boundary condition. For the full derivation and explanation of the coefficients, we refer to Cuccia et al. [3].

## 3. Results and Discussion

### 3.1 Validation on Homogeneous Phantoms

To first validate the laparoscopic SFDI system, measurements were performed on homogeneous intralipid phantoms of known optical properties. After acquiring SFDI data on all phantoms, images were averaged and the mean percent error was calculated. Additionally, the coefficient of determination (r^2^) between measured and true values was calculated to confirm precision and linear sensitivity to varying optical properties. In Fig. 4, we demonstrate representative plots and calculations with measured data. In the absorption variation experiment, recovered versus expected absorption shows linearity. When absorption is very high, a slight overestimation trend is observed independent of calibration choice. The experiment’s recovered reduced scattering values show less than 6% deviation from the expected value in all cases. Similar linearity is observed in the reduced scattering variation experiment. Absorption values in this case demonstrate less than 10% deviation from the expected value. Our laparoscopy SFDI system shows agreement that has been reported earlier [14,41,42].

**Fig. 1.**
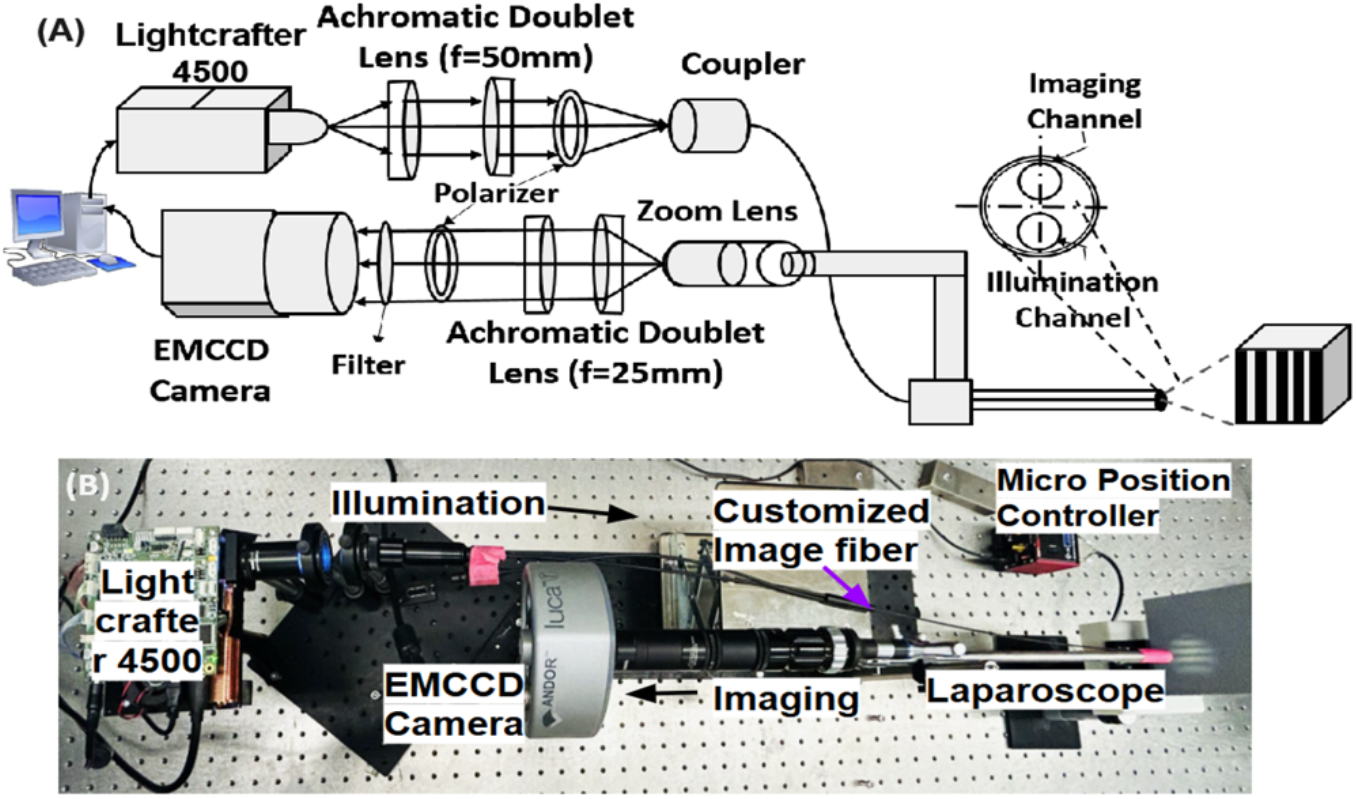
a) Schematic of wide-field laparoscopic system used to acquire all phantom data with ray diagram of structured light projection optics. (b) LSFDI prototype.

**Fig. 2.**
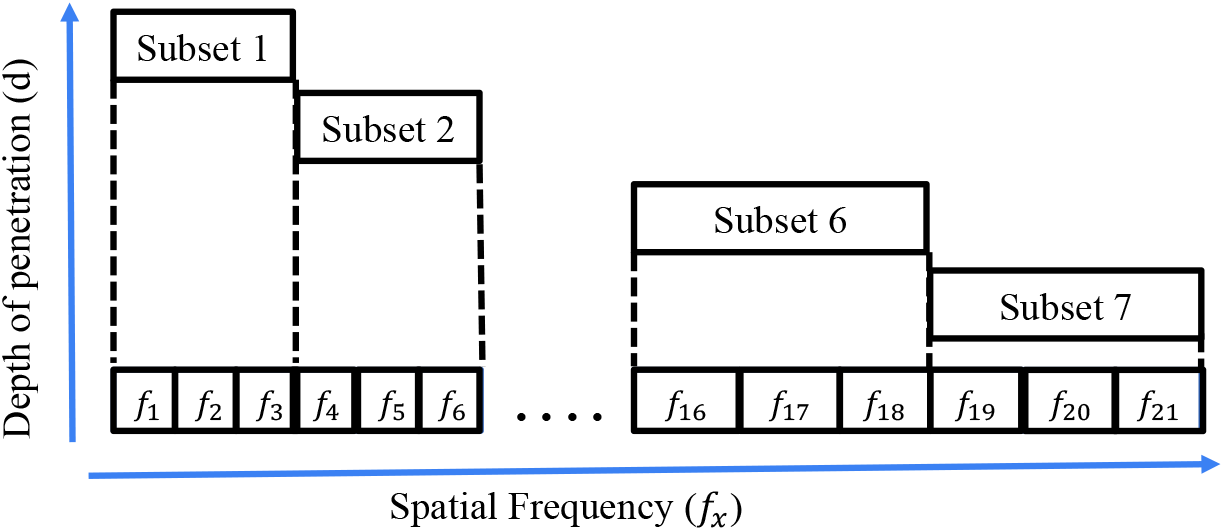
The dataset of 21 spatial frequencies (*f*_*x*_) was divided into 6 smaller, sequential sub-sets, each containing 3 *f*_*x*_ values. Each subset is associated with a different penetration depth (δ), which is estimated based on the average spatial frequency of the sub-set, where a lower *f*_*x*_ corresponds to a greater δ.

**Fig. 3.**
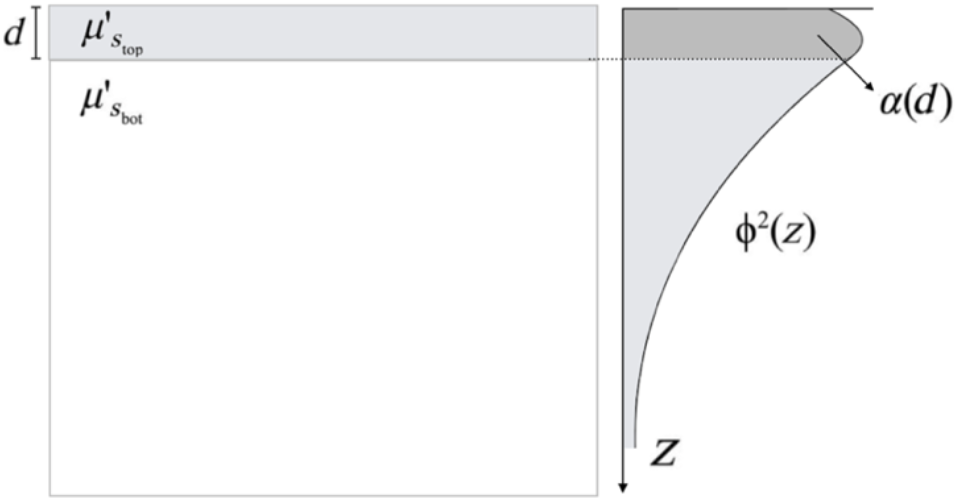
A 2D illustration of the two-layer scattering model. A homogeneous thin layer of thickness d and scattering coefficient 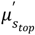 lies atop a semi-infinite, also homogeneous layer with scattering coefficient 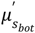. The light fluence *ϕ* is shown to decay exponentially with depth (z-direction) from the surface. The weighting coefficient *α(d)* quantifies the partial contribution of the fluence squared (*ϕ*^*2*^) within the top layer. This is calculated by integrating *ϕ*^2^ from the surface to depth d and normalizing it by the total integral of *ϕ*^*2*^ over all depths.

**Fig. 4.**
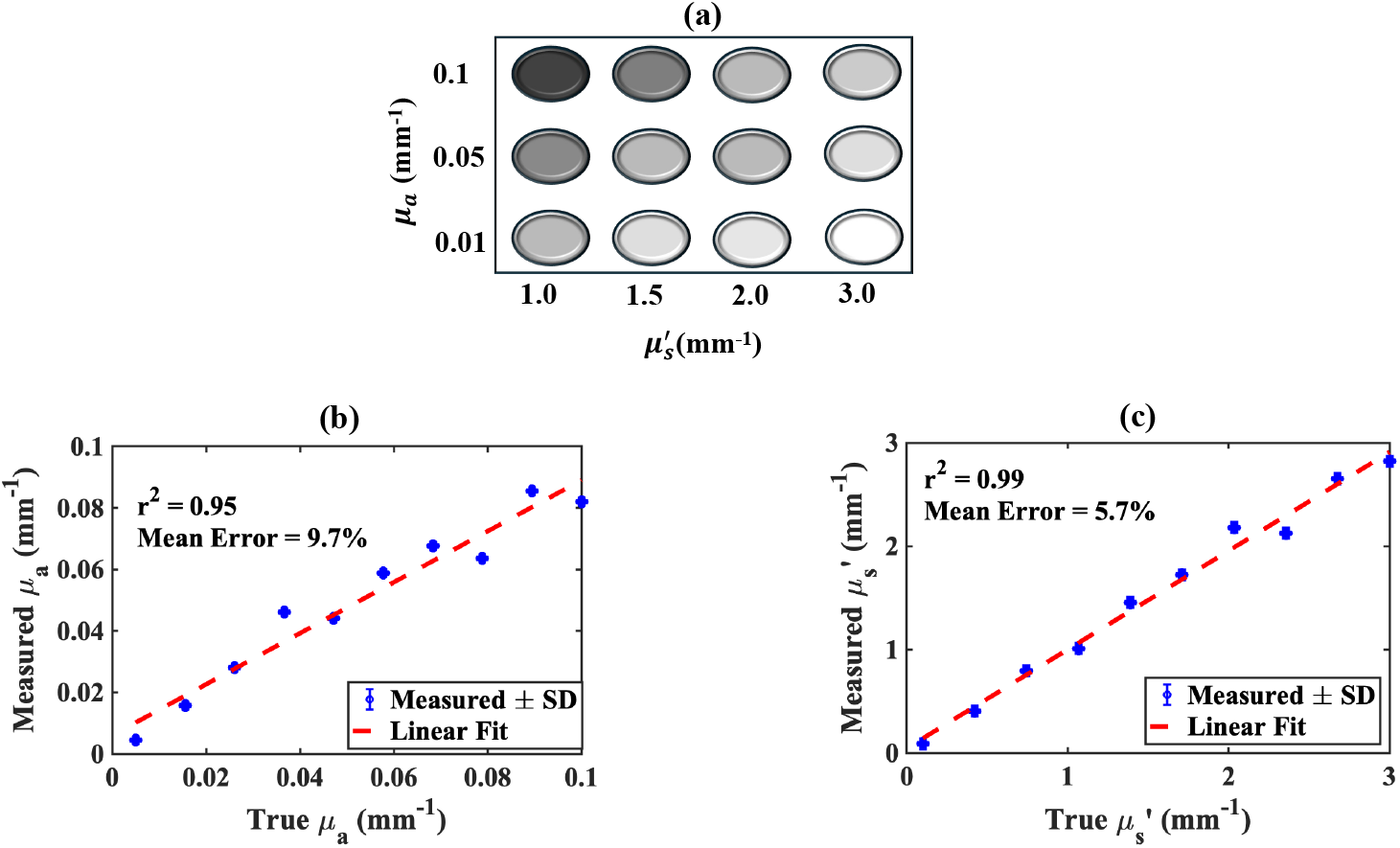
Validation of the laparoscopy SFDI system with different phantoms. (a) Sample of phantoms used to validate the system, (b) Mean percentage error and correlation of measured vs true absorption coefficient and (c) Mean percentage error and correlation of measured vs true reduced scattering coefficient. Error bars and correlation line show how the true and measured optical properties are different from each other.

### 3.2 Silicone/Silicone Phantom Configuration

Fig. 5 presents the measured and modeled scattering coefficients 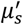 for the silicone/silicone model using the three different modeling techniques described in 2.4. The figure focuses on a single spectral band (656 nm), and the data is displayed as a function of spatial frequency with the average spatial frequency within each of the six frequency subsets on the horizontal axis. For reference, the scattering coefficients of the top and bottom layers are shown, both assumed as constant across spatial frequencies. These reference values were obtained from measurements on thick, homogeneous phantoms using the same SFDI system.

**Fig. 5.**
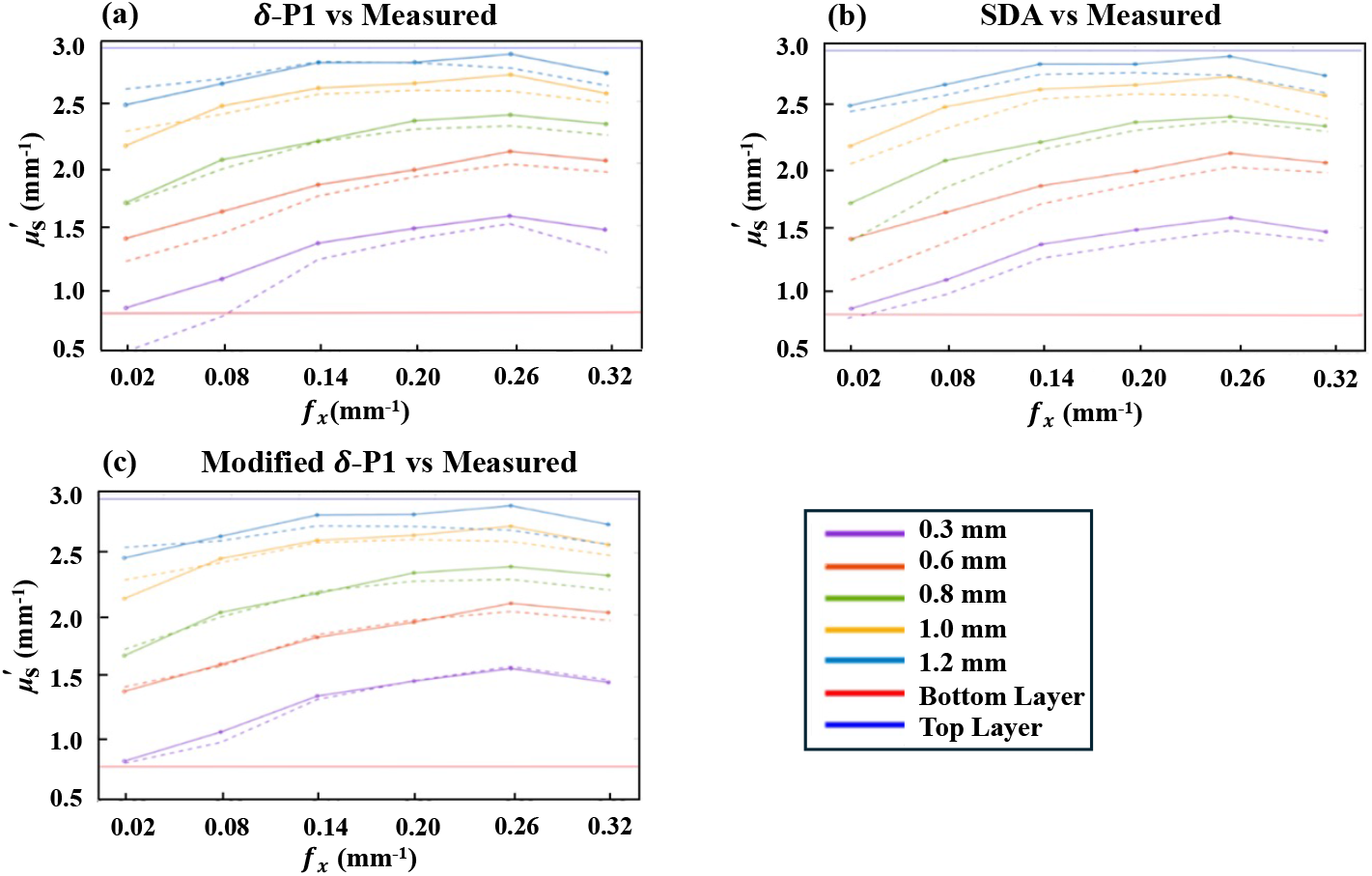
Silicone/Silicone Model. Measured 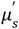 of multi-layer model at 6 different spatial frequency subsets (solid lines) plotted against (a) δ-P1 approximation, (b) SDA and (c) a modified δ-P1 approximation (dashed lines) on silicone phantoms of different thicknesses but identical optical properties.

The measured values for the two-layer phantoms lie between the top and bottom phantom 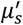 aligning with our theoretical model where the response is a combination of both layers. As the thickness of the upper layer increases, the scattering values shift closer to the top layer as expected, and as the average value of the analyzed projected frequency increases, 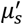 increases as well. This is consistent with SFDI behavior where higher spatial frequencies correspond to shallower light penetration, thereby enhancing the influence of the top layer. This relationship does not appear linear, but rather plateaus or “saturates” as either projected frequency or layer thickness increases. This saturation effect, occurs notably from 0.14 to 0.26 fx and is attributed to the light’s penetration depth becoming smaller than the thickness of the top layer, effectively limiting the measurement to a single-layer response.

### 3.3 Silicone/Intralipid Phantom Configuration

Fig. 6 displays the silicone/intralipid model and shows a similar trend as the silicone/silicone model. As seen in Fig. 5, the measured 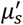 values increase with different frequency subsets and top layer thicknesses. However, this saturation effect is less pronounced in the silicone/intralipid model, where at thicker layers the plateau occurs at higher frequencies (0.3 fx) compared to the silicone/silicone model (0.14 fx).

**Fig. 6.**
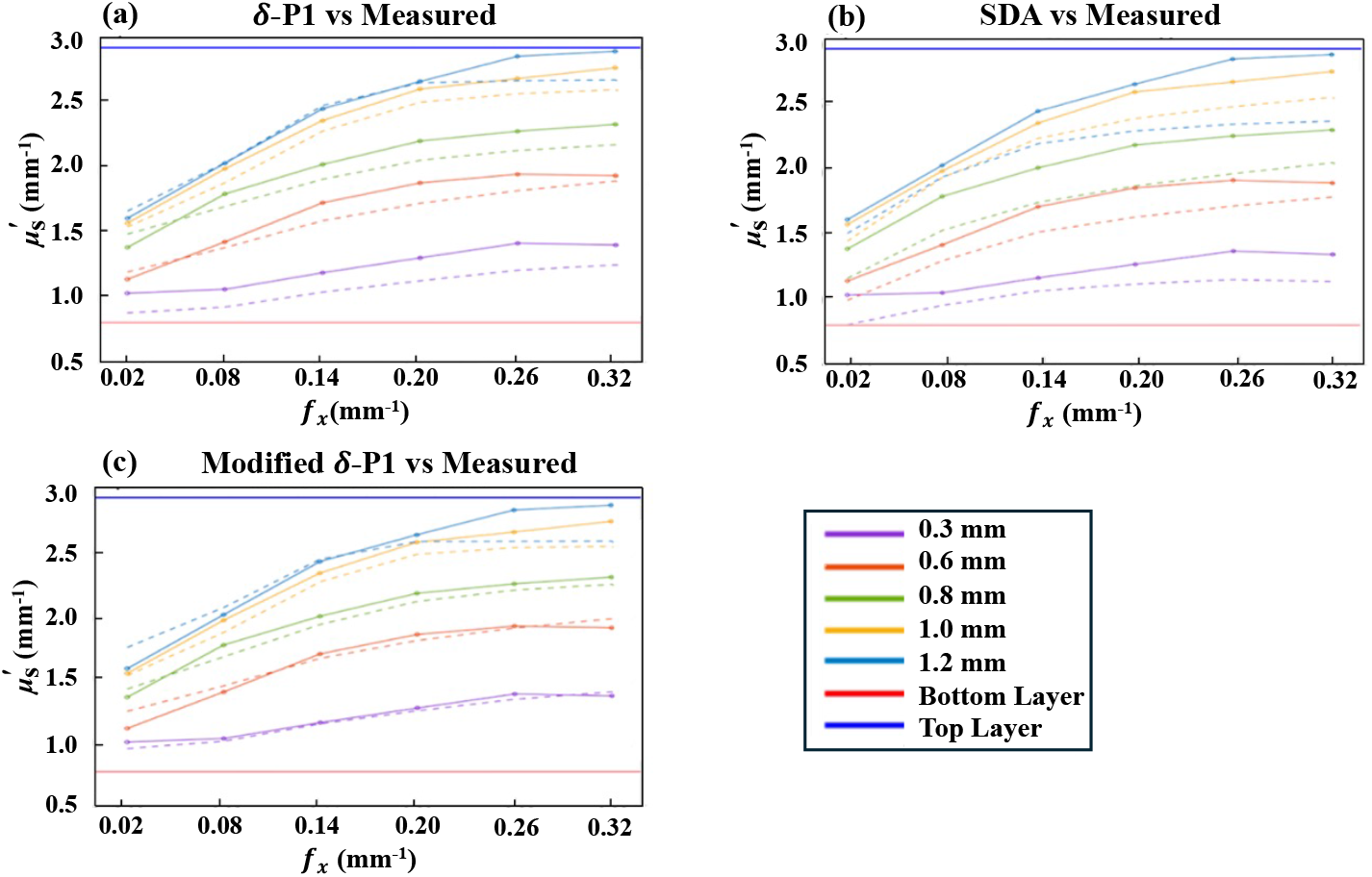
Silicone/Intralipid Model. Measured 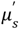 of multi-layer model at 6 different spatial frequency subsets (solid lines) plotted against (a) δ-P1 approximation, (b) SDA and (c) a modified δ-P1 approximation (dashed lines). Models were modified via equation 3 to account for index of refraction changes at the silicone/intralipid interface.

There are a few differences between the silicone/silicone model and the silicone/intralipid model that can be attributed to the different index of refraction (*n*) changes between the two systems. In the silicone/intralipid model, we see lower frequency 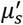 values significantly less than the silicone/silicone model, which might be due to photons being “trapped” at the silicone/intralipid boundary. Once in the intralipid, photons are likely to have undergone many scattering events and are more likely to be beneath the angle of incidence required to pass through back to the top layer, thus increasing absorption in the bottom layer. Additionally, while care was made to remove visible air pockets in both models, there may be microscopic air gaps between the silicone/silicone layers not present in the silicone/intralipid model. With air having *n* = 1.0 and intralipid having *n ≈* 1.33, any small air pockets between the layers would trap more photons within the top layer, inflating the recorded 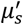.

Fluence modeling trends for the SDA, δ-P1, and Modified δ-P1 models follow their expected limitations. The SDA underestimates top-layer contributions, particularly at low frequencies, due to its reliance on long-path diffusion assumptions (i.e., distance from source ≫1/*μ*.). The original δ-P1 model reduces bottom-layer overestimation at low frequencies by accounting for short-path events, resulting in improved 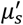 estimation. The Modified δ-P1 model performs best for very thin top layers but increasingly overestimates top-layer influence as thickness increases.

### 3.4 Evaluating Model Accuracy

To objectively evaluate and compare the performance of the two-layered 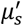 models, the Root Mean Square Percentage Error (RMSPE) was calculated for each model across all phantom thicknesses. Using the measurements detailed in Sec. 2.3, the calculations were performed via the following equation:

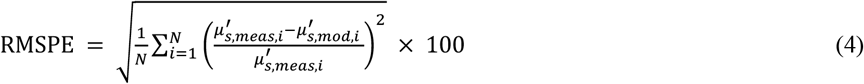

In this expression: N represents the number of average spatial frequencies in the dataset 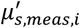 is the measured scattering coefficient and 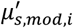 is the scattering coefficient derived via Eq. (1). The resulting performance data is summarized in Fig. 7, which charts the RMSPE between all three models on the silicone/silicone and silicone/intralipid phantom configurations. Additionally, Fig. 8 compares the RMSPE of each model between phantom configurations to demonstrate changes in model behavior in the context of a refractive index change.

**Fig. 7.**
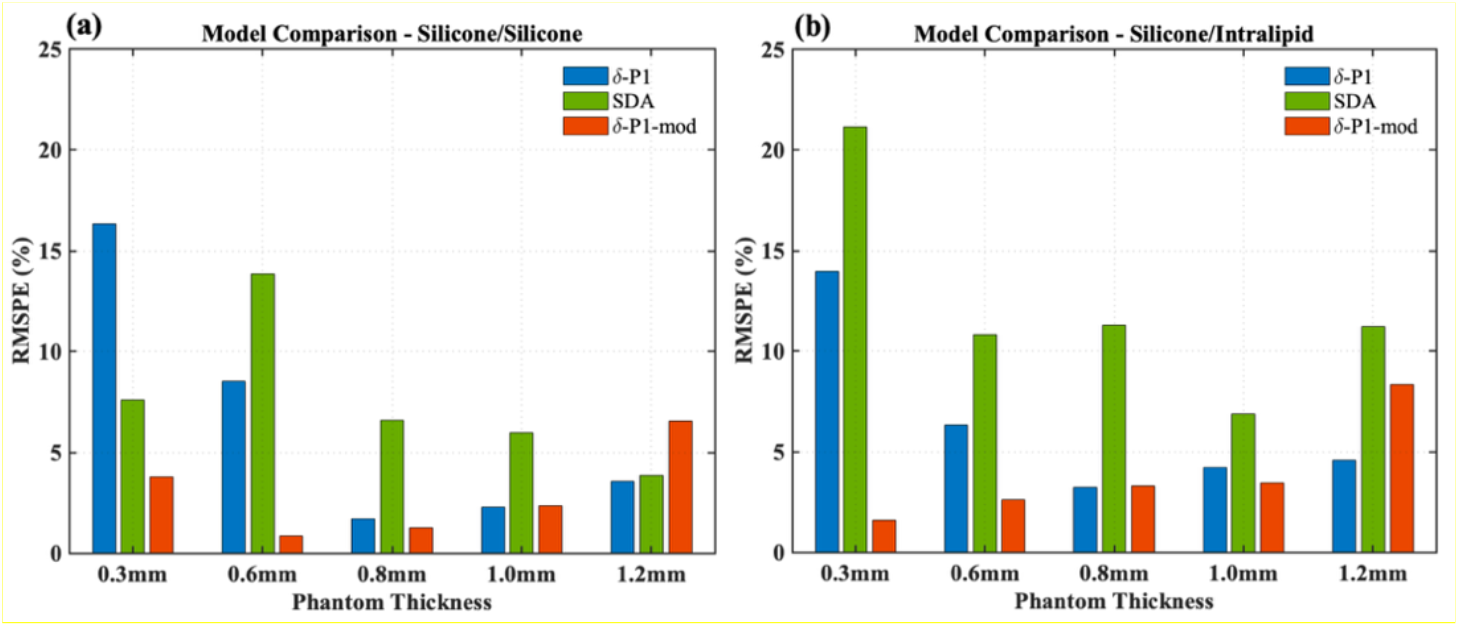
To evaluate the accuracy of the three fluence-based two-layer models, the RMSPE was calculated based on the deviation between the modeled scattering coefficients and the experimental measurements. Within each figure, the RMSPE is displayed for all three models across the five tested top-layer thicknesses, with (a) showing values for the silicone/silicone and (b) the silicone/intralipid phantom configurations.

**Fig. 8.**
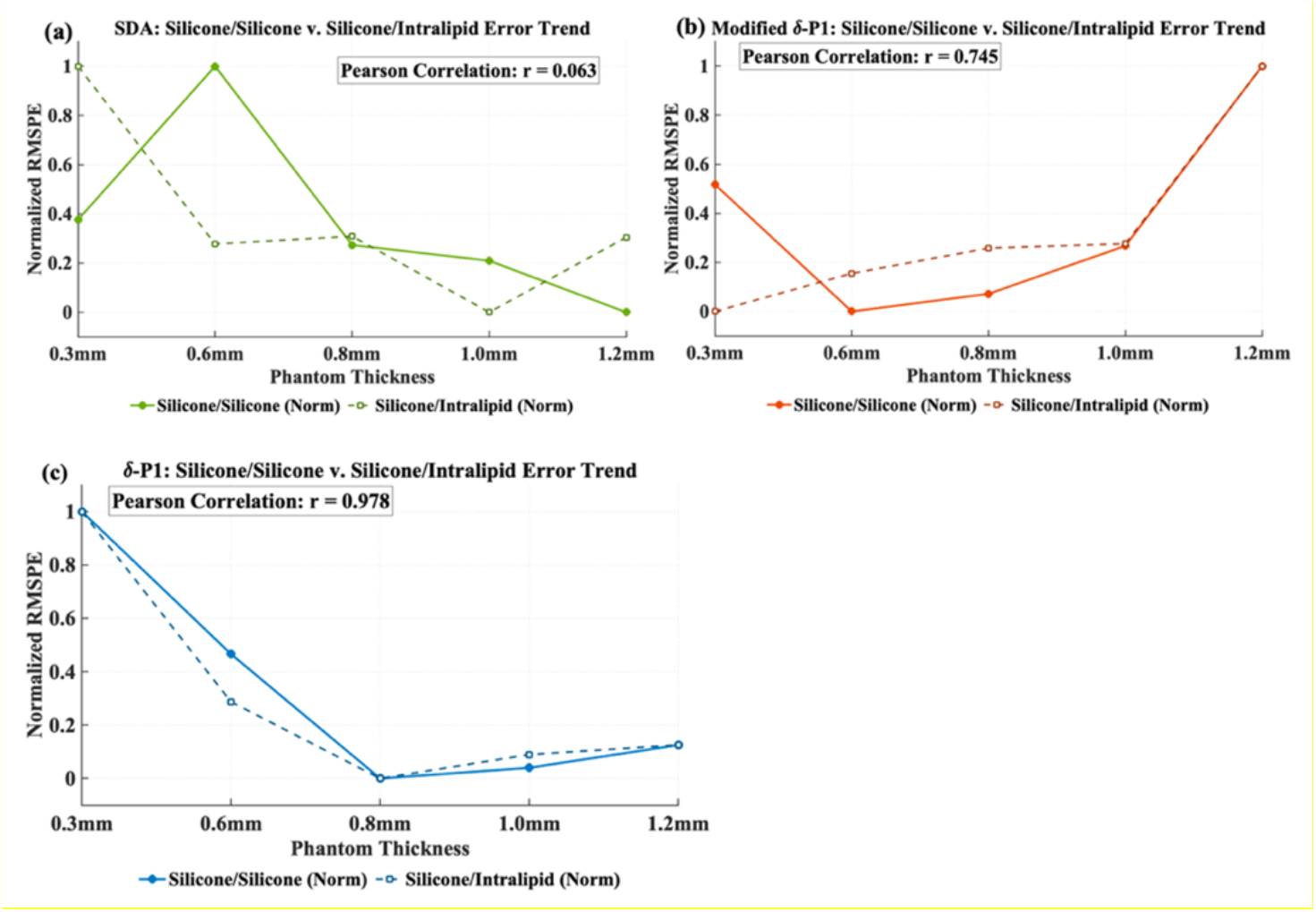
Comparison of RMSPE trends across varying top-layer thicknesses for silicone/silicone and silicone/intralipid layered phantoms with calculated Pearson correlation coefficient. The plots illustrate changes due to index of refraction change between the three different modeling techniques: (a) SDA model exhibits poor correlation between the two phantom types r = 0.063, (b) Modified δ-P1 shows improved trend consistency with r = 0.745 and (c) the δ-P1 model is most consistent across phantom configurations with r = 0.978.

**Fig. 9.**
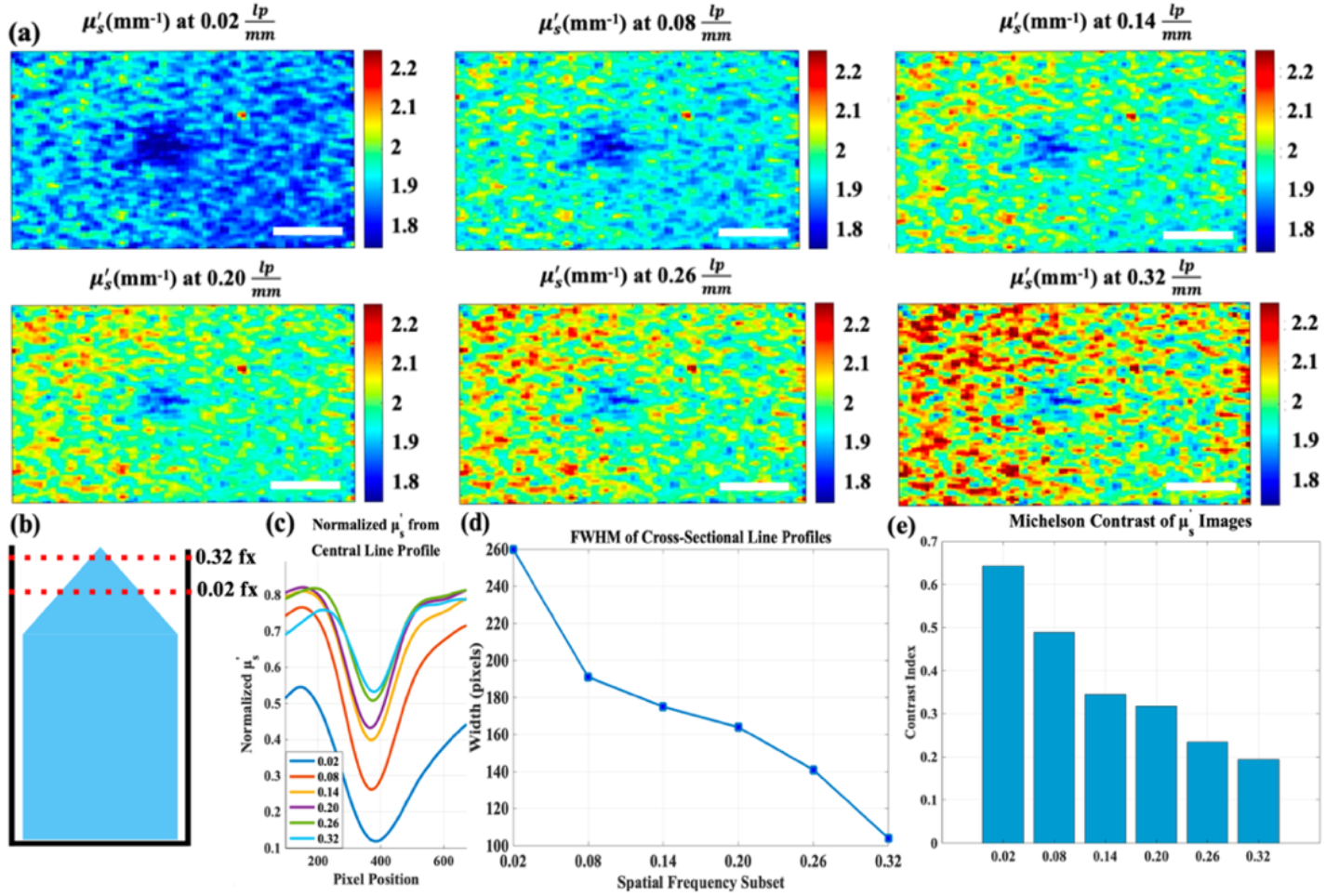
A cone shaped, low 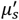 phantom was placed in a cup filled to the brim with high 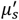 intralipid (b) and imaged at increasing spatial frequency subsets to visually demonstrate SFDI depth sensitivity. (a) shows an inverse relationship between spatial frequency subset and the contribution of the cone’s 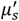(c) show smoothed horizontal line profiles extracted from the spatial center of each sample, (d) and (e) show derived FWHM and contrast index respectively. Scale bar = 8mm.

Furthermore, introducing a refractive index mismatch differentiates model behavior. The SDA exhibits substantial sensitivity to phantom type despite identical 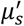, as Fresnel reflections and abrupt trajectory changes violate diffusion assumptions. In contrast, both δ-P1 models remain more consistent across refractive index transitions by incorporating short-path photon corrections, making them more robust to interface-induced optical disruptions.

### 3.5 Cone Phantom for 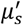 Depth Gradient

Fig. 9 gives a visual example of how the differing penetration depth of the projected sinusoidal frequencies affects the derived optical properties of a heterogeneous phantom. Low frequencies see a much larger area containing the optical properties of the submerged silicone phantom, while higher frequencies see a smaller area with less penetration. Notably, the derived 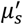 of the intralipid phantom increases along with the projected pattern frequency, which is expected due to high-frequency data being less sensitive to the absorption contrast within the intralipid phantom [3]. The line profiles in Fig. 9c show a drop in 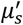 at the point where the tip of the cone is under the surface of the intralipid. The width of this drop and its relative contrast across the entire image is largest at low frequencies and smallest at high frequencies, as shown by Fig. 9d, e. This “SFDI sectioning” [16] in a laparoscopic system can be used to detect cancerous tissue with abnormal optical properties hidden by surface layers.

The reduced scattering coefficient serves as the main parameter of analysis for depth resolution due to the relatively limited likelihood of an absorption event occurring at the probed volumes. At higher spatial frequencies, the photon path length is sufficiently short that the probability of absorption is negligible relative to multiple scattering events, a likelihood compounded by the lack of significant vascularization or pigment densities within superficial peritoneal layers. By isolating heterogenous scattering contrast, our model can avoid the potential for confounding absorption effects of deeper, more vascularized tissue [43].

This study focused on applying depth-resolving SFDI techniques to data acquired across multiple spatial frequencies within a laparoscopic system. Although observed 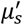 trends match theoretical expectations, current models require prior knowledge of tissue geometry and optics, limiting intraoperative applicability. Future work will therefore develop an iterative inverse solver to jointly estimate layer scattering and thickness directly from raw multi-frequency measurements and extend the method to multi-wavelength 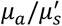 mapping at fluorescence excitation and emission bands. These advances will enable depth-aware fluorescence attenuation correction and light-dose modeling to support image-guided CPT treatment planning and monitoring, including quantitative mapping of PoP photobleaching and light-triggered doxorubicin release. Importantly, depth tuning via spatial frequency provides a direct bridge to CPT: lower frequencies probe deeper volumes relevant to treatment-light penetration, while higher frequencies emphasize superficial layers that dominate fluorescence escape.

## 4. Conclusion

This study demonstrates depth-sensitive optical property characterization with a laparoscopic SFDI system by partitioning the measured spatial-frequency content into multi-frequency subsets with distinct effective penetration depths. Using two-layer tissue-mimicking phantoms, we show that the recovered 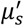 transitions systematically between bottom-and top-layer values as the superficial-layer thickness and spatial frequency increase, consistent with depth-dependent sensitivity predicted by two-layer scattering models. Among the evaluated analytical models, the δ-P1 variants generally outperformed the standard diffusion approximation, with the modified δ-P1 model providing the best agreement for very thin (≈0.1-0.2 mm) superficial layers relevant to epithelial tissue. We further evaluated refractive-index mismatch to emulate additional tissue heterogeneity and observed similar depth-dependent trends. Together, these results establish a practical pathway toward depth-aware 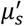 estimation in laparoscopic imaging, which will be leveraged in future work to improve depth-weighted quantitative fluorescence correction and light dosimetry for fluorescence-guided drug delivery and chemophototherapy monitoring in ovarian cancer.

## Acknowledgments

This research was supported by the National Institutes of Health/National Cancer Institute under grant number R01CA243164 (5R01CA243164-06). ChatGPT 5.2 was used to assist with grammar and language editing during manuscript preparation.

## Data availability

Data and materials supporting the findings of this study are available from the corresponding author upon reasonable request.

